# Badger territoriality maintained despite disturbance of major road construction

**DOI:** 10.1101/2020.11.06.370957

**Authors:** Aoibheann Gaughran, Enda Mullen, Teresa MacWhite, Peter Maher, David J. Kelly, Ruth Kelly, Margaret Good, Nicola M. Marples

## Abstract

Road ecology has traditionally focused on the impact of in-situ and functional roads on wildlife. However, road construction also poses a major, yet understudied, threat and any implications for key aspects of animal behaviour are unknown. There are also concerns that environmental disturbances, including major roadworks, can disrupt badger territoriality, promoting the spread of tuberculosis to cattle. To address these knowledge gaps, the ranging behaviour of a medium-density Irish badger population was monitored, using GPS-tracking collars, before, during and after a major road realignment project that bisected the study area. We estimated badgers’ home range sizes, nightly distances travelled and the distance and frequency of extra-territorial excursions during each phase of the study and quantified any changes to these parameters. We show that roadworks had a very limited effect on ranging behaviour. A small increase in nightly distance during the roadworks did not translate into an increase in home range size, nor an increase in the distance or frequency of extra-territorial excursions during the roadworks. In addition, suitable mitigation measures to prevent badger deaths appeared to ensure that normal patterns of ranging behaviour continued once the new road was in place. Our analysis supports the view that road construction did not cause badgers to change their ranging behaviour in ways likely to increase the spread of tuberculosis.

## INTRODUCTION

Studies on the impact of roads on animals have tended to focus on their impact on population density; barriers to, or facilitators of, movement; habitat loss and fragmentation; mortality; increase or decrease of food availability; and avoidance behaviour once roads are already constructed and in-situ *e.g.* (Carr *et al*. 2002; Coffin 2007; Benítez-López *et al*. 2010). However, there have been few studies that have specifically examined the impact of the process of road construction on animals (Torres *et al*. 2011). This lack is surprising because the building of roads could have an extremely high disturbance intensity (Kohn *et al*. 1999; Klar *et al*. 2009; Lesmerises *et al*. 2013), including vegetation clearance, rock-breaking, major excavation and earth-moving activity. This study provides insights into the impacts of road construction on these key elements of animal behaviour using a large GPS-tracking dataset of European badger *(Meles meles).* Most of the literature pertaining to the disruptive effects of roads on mustelid ecology *(e.g.* Clevenger 2013), and specifically badger ecology, consider barriers to dispersal, mortality due to road traffic accidents (RTAs) and the mitigation of these effects (Davies *et al*. 1987; Van der Zee *et al*. 1992b; Clarke *et al*. 1998; Dekker and Bekker 2010). Little is known about the impact of road construction on badger behaviours such as movement and territoriality.

The badger makes an appropriate study species because it is subject to protection under Annex III of the Bern Convention (1979) under which activities capable of causing local disappearance of, or serious disturbance to, populations are prohibited. However, in some European countries, badgers are also subject to control. In both Ireland and the UK, badgers have been implicated in the spread of *Mycobacterium bovis* to cattle, and act as a wildlife reservoir for bovine tuberculosis (bTB) (Gormley and Costello 2003; Murphy *et al*. 2010; Corner *et al*. 2011; Godfray *et al*. 2013). TB transmission between badgers and between badgers and cattle is highly complex and multi-factorial, involving many other factors such as cattle-to-cattle transmission, movement of infected livestock between herds, biosecurity and the poor sensitivity of TB testing in cattle (O’Connor *et al*. 2012; Allen *et al*. 2018). Nonetheless, in order to understand the dynamics of a disease in a wildlife reservoir, and to control it successfully, as complete a picture as possible of the ranging behaviour of the carrier species is required (Conner and Miller 2004).

The organisation of badgers into territorial social groups appears to limit the spread of TB because it lowers disease transmission rates between groups (Cheeseman *et al*. 1988; Delahay *et al*. 2000; Davis *et al*. 2015). However, badger movements into and out of neighbouring social groups are associated with increased prevalence of TB in those groups (Rogers *et al*. 1998; Riordan *et al*. 2011). Therefore, the ranging behaviour of badgers is of direct importance to the transmission of TB infection both between individual badgers (Weber *et al*. 2013; O’Mahony 2015) and between badgers and cattle (Martin *et al*. 1997; Eves 1999; Griffin *et al*. 2005b; Woodroffe *et al*. 2009; Mullen *et al*. 2015).

From studies of disease control, there is evidence for both positive and negative effects of badger culling on bTB breakdown rates. While Irish studies have found that lowering population density reduces TB transmission (Eves 1999; Griffin *et al*. 2005b, a; Olea-Popelka *et al*. 2009), studies in the UK have found that culling may produce a ‘perturbation effect’, *i.e.* it may disrupt badger ranging behaviour such that TB transmission to cattle increases resulting in an increase in herd breakdowns (Cheeseman *et al*. 1993; Tuyttens *et al*. 2000a, b; Donnelly *et al*. 2006, 2007; Carter *et al*. 2007; Jenkins *et al*. 2008). These disruptions to ranging behaviour include rapid immigration into culled areas, increased social group range size, increased social group range overlap, increased inter-group movements and increased ranging of individuals (Carter *et al*. 2007), all of which result in higher TB transmission rates among badgers and between badgers and cattle (Woodroffe *et al*. 2006; Donnelly *et al*. 2007; McDonald *et al*. 2008; Godfray *et al*. 2013). In contrast to the perturbation effects found in the UK, Irish studies have reported sustained decreases in the rates of bTB breakdowns as a result of culling (Eves 1999; Griffin *et al*. 2005b, a; Olea-Popelka *et al*. 2009), despite some evidence for disruption of ranging behaviour (O’Corry-Crowe *et al*. 1996; Costello *et al*. 2006). Other human activities have the potential to disturb badger ranging behaviour and consequently influence the rate of disease transmission. For example, density reduction due to persecution has been found to increase the distance badgers travel, and to reduce territoriality (Sadlier and Montgomery 2004; Sleeman and Mulcahy 2005) and has been associated with increased risk of TB in cattle and the persistence of TB hotspots (Wright *et al*. 2015). It has been suggested that environmental disturbances, such as clear felling of forests (Blumstein 2010; Mullen *et al.,* 2019.) and landscape changes that occur as road density increases (Van der Zee *et al*. 1992a), may also impact the population density and ranging behaviour of badgers. Disruptive events like these may also have implications for TB transmission among badgers and between badgers and cattle. Recently, the Irish media have reported concerns that the construction of major roads may increase bTB breakdowns resulting from an associated increase in movement by badgers *(e.g.* Hubert 2016; Ó Liatháin 2020).

Ranging behaviour can be described by a variety of movement metrics, including home range size (HR), nightly distance moved (ND), the distance of extra-territorial excursions (ETEs) and the frequency of ETEs (fETE). The distances badgers move each night vary considerably, from an average of 7km per night at low population densities (Kowalczyk *et al*. 2006) to an average of less than 1km (849m) per night (Woodroffe *et al*. 2016) at higher population densities. Within territories, the sizes of home ranges vary seasonally, being smallest in winter (Kowalczyk *et al*. 2003; Do Linh San *et al*. 2007; Palphramand *et al*. 2007) and largest in summer (Revilla and Palomares 2002; Kowalczyk *et al*. 2003; Kauhala *et al*. 2006; Woodroffe *et al*. 2016). HR also varies with population density, from 22km^2^ in low density populations (Kowalczyk *et al*. 2000) to 0.26km^2^ in very high-density populations (Chris Newman, pers. comm.) The majority of Irish populations (76%) are of medium density, averaging 1.4 badgers km^-2^ (Table S1). A large-scale trapping study in Ireland (Byrne *et al*. 2014) found that movements of >1km, which would take a badger outside its territory, accounted for 57% of all movements detected. Irish badgers have been recorded making such ETEs in all seasons, some of which may be up to 8km from their home territory (Sleeman 1992; Sleeman and Mulcahy 1993).

If roadworks disrupt badger ranging behaviour, we would expect to see changes in any or all ranging parameters. The distance moved in a night (ND) might increase during the construction phase, when disturbance is greatest (see Carter *et al*. 2007). Such an increase in ND could result in a corresponding increase in HR and/or an increase in ETE distance and frequency, as badgers seek refuge from the ecological, acoustic and seismic disturbance of roadworks. If extreme, these disturbances might lead to a breakdown in territoriality in the area around the roadworks.

The present study monitored the ranging behaviour of a medium-density population of badgers over 6.5 years: before, during and after a major road upgrade. This allowed us to assess how road-building activity affected the ranging behaviour of badgers, and how rapidly recovery, if any, occurred. We considered the before/during/after comparisons with respect to their ecological importance, as well as their statistical significance. Such a long-term, detailed study of badger ranging behaviour using GPS tracking technology has never been conducted before.

## MATERIALS AND METHODS

The study was conducted in mixed farmland in Co. Wicklow, Ireland (52.924130, −6.117960). A description of the study area and details of the trapping and handling of badgers are given in Gaughran *et al*. (2018). Over the course of the study we trapped 139 badgers, 80 of which wore GPS tracking collars. The population density in the study area was 1.8 badgers/km^2^ and remained stable over the 6.5-year study period (Gaughran *et al*. 2018). The average distance between main setts was 1.3km. The road upgrade involved building a new 16km section of motorway (M11) alongside the original national road (N11) (MacWhite *et al*. 2013). The area where roadworks occurred was long and narrow, on average 100m wide (max. 400m) and 16km in length (Fig. S1). Wildlife underpasses were installed under the M11 at appropriate locations following surveys to locate badger paths and crossings, the location of RTAs and the examination of GPS data from the ‘before’ phase. The entire 16km of new motorway was lined with badger-proof fencing which effectively guided wildlife into underpasses. The mitigation measures were checked by the National Parks and Wildlife Service (NPWS) to ensure functionality and compliance to statutory requirements.

Ethical approval for the project was granted by Trinity College Dublin’s Animal Research Ethics Committee (Project No. 290516) and the Health Products Regulatory Authority (Project No. 7024754). Badgers were captured under licences (NPWS Nos. 101/2009, 04/2010, 13/2010, C123/2010, 03/2011, C040/2011, C03/2013, C005/2013 and C001/2015) as required by the Wildlife Act, 1976. Both cage traps and stopped-restraints conformed to national legislation for humane trapping defined in the Wildlife Act, 1976, Regulations 2003 (S.I. 620 of 2003) (MacWhite *et al*. 2013).

### Data Collection & Management

Badgers were tracked using Tellus Light GPS collars (Followit Wildlife, Lindsberg, Sweden). Data collection began in April 2010 and GPS tracking continued until October 2016, when all collars were removed. We aimed to capture as many badgers as possible within each social group. Captured badgers were anaesthetised in-cage by veterinary practitioners at each trapping event (Gaughran *et al*. 2018). Badgers were sexed and had their teeth examined and photographed. Age was determined by dentition (Hancox 1988; da Silva and Macdonald 1989) and general appearance of each badger. Irish badgers give birth from the first week in February to early March with the majority of cubs born in mid-February (Corner *et al*. 2015). For convenience in age classification, the date of birth was assumed to be the 1^st^ of February for all badgers. Age cohorts were defined as follows: cub (a badger in its first year); young adult (a yearling or a 2-year old); older adult (a 3- or a 4-year old) and aged adult (badgers ≥5 years old).

A social group was defined as the group of badgers which were regularly trapped at the same main sett and whose home ranges overlapped during the time-period in question (Macdonald *et al*. 2008; Woodroffe *et al*. 2016). Thus, badgers were assigned to a social group based on their most frequent trapping location and, if collared, their GPS tracking data. Badgers were categorised into one of three ranging status categories - traditional rangers, dispersers or super-rangers - on the basis of their behaviour at the time that the GPS location was recorded. Dispersers (confirmed retrospectively) were badgers in the process of moving permanently from one social group to another social group (Gaughran *et al*. 2019). Super-rangers were those badgers that ranged within an extended territory consisting of their original/natal territory and some, or all, of an adjacent territory or territories (Gaughran *et al*. 2018). The vast majority of the population were traditional rangers, accounting for ≥80% of movement observations. Both super-rangers and dispersers were excluded from the analyses presented here, as the roadworks were found to have had no impact on their ranging behaviour (Gaughran 2018).

Social groups were categorised as adjacent to the roadworks if they shared a border with the N11/M11 and associated roadworks. Those social groups that were further away were categorised as non-adjacent. The timing of the roadworks was split into three phases: ‘before’ (April 2010-August 2013, 41 months), ‘during’ (September 2013 - June 2015, 22 months) and ‘after’ (July 2015 - August 2016, 14 months) the construction of the road. During the ‘before’ period the area where the new motorway was to be located (Fig. S1, Fig. 1 a) was cleared of trees, hedgerows and grass/crops and became mostly scrub land. Despite being fenced, it was completely accessible to and regularly used by badgers. In the ‘during’ phase, the scrub within this area was completely cleared, leaving exposed earth. The roadworks involved rock-breaking, major excavation and earth-moving activity within the construction zone (Fig. 1b). Continuous badger-proof fencing was installed only at the very late stages of the ‘during’ phase. In the ‘after’ phase, badger proof-fencing and underpasses were fully operational (Fig. 1c). Badgers did not have access to the area around the M11 and its associated access roads. Underpasses were inspected for accessibility and their use by badgers (and other wildlife) confirmed through the use of camera-traps.

**Fig. 1.**
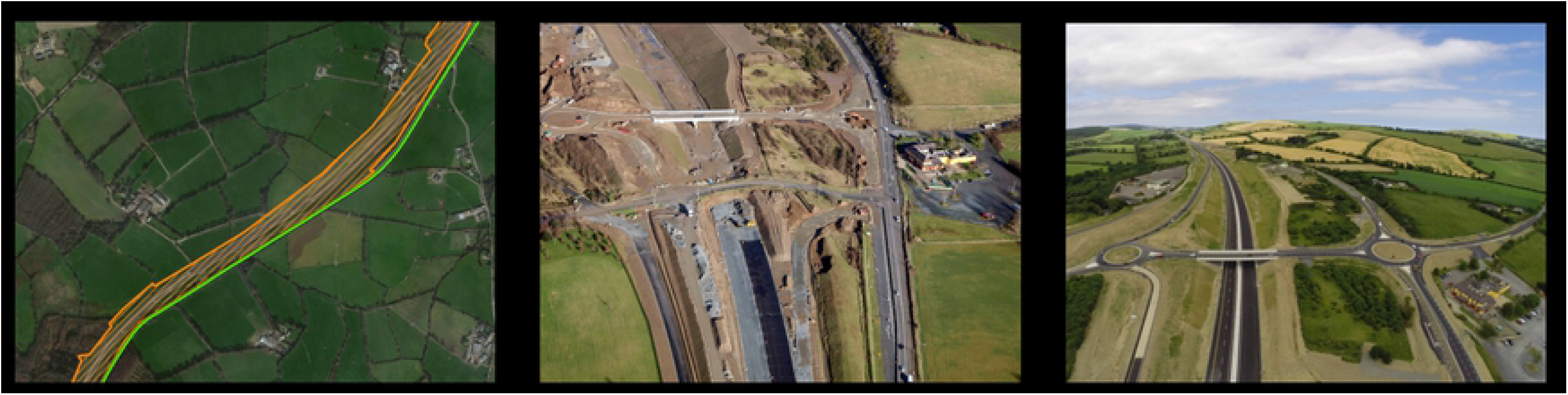
Roadworks phases. **a) ‘Before’** satellite image of part of the study area before the roadworks. The hatched orange area indicates the location of the M11 motorway construction zone running alongside the N11 road. The N11 is represented by the green line. **b) ‘During’** - aerial photograph of part of the study area taken during the roadworks revealing the extent of the change to the habitat. The N11 can be seen to the right-hand side of the photograph. **c) ‘After’** - aerial photograph of part of the study area taken upon completion of the roadworks. The motorway and access roads were completely enclosed in continuous badger-proof fencing.

GPS collars were programmed to record four locations a night, at 21:00, 23:00, 01:00 and 02:00, for each collared badger. GPS locations from the Followit website were converted from the global geodetic system WGS-84 to the Irish Grid projection system using Grid InQuest II (Ordinance Survey Ireland). Data were visualised by mapping the GPS locations onto the World Imagery basemap using the coordinate system TM_75 Irish Grid in ArcMap (ArcGIS version 10.4.1). The study area was digitised by creating a polygon shapefile, which traced the landscape features in the study area such as field boundaries, roads, rivers, forestry, and farmyards. This resulted in a dataset of 81,925 GPS records from April 2010 to August 2016 (77 months) from 80 different individuals.

To calculate ND, for each collared badger, nightly trajectories were created using the package *adehabitatLT* (Calenge 2011) in R Version 3.4.0 (R Core Team 2016), in which a ‘burst’ was defined as a single night’s GPS locations, and the total distance travelled each night was extracted in meters (m). Two or more GPS locations were required to calculate ND, but, because collars could not transmit when the badger was below ground, and badgers did not always emerge from their setts for the full night, badgers did not always send the four scheduled GPS locations in a night. Therefore, our analysis only considered data from those nights on which badgers successfully recorded two or more GPS locations. Our calculations of how far a badger travelled in a night will inevitably be under-estimations for two reasons. Although badgers tend to follow linear features in their landscape (O’Brien *et al*. 2015), their routes can be quite tortuous, particularly when foraging (Loureiro *et al*. 2007). Our analysis was limited to considering straight line distances between the four GPS locations. Secondly, the dataset is lacking information regarding how far a badger travelled both before and after the first and last GPS locations were recorded. However, as the methods were identical throughout the study, this should not influence our ability to make statistical comparisons between time-periods or groups of badgers.

To generate the HR dataset, monthly home range size was estimated by calculating 95% minimum convex polygons (MCPs) for each badger in each calendar month of each year of the study using the R package *adehabitatHR* (Calenge 2006). MCPs could only be calculated when there were >5 records for a badger in a given month. We chose to use 95% MCP as a proxy for home range size as it is comparable with the majority of existing badger studies (Elliott *et al*. 2015).

ETEs are journeys made by badgers beyond the boundaries of their social group’s territory into neighbouring territories or beyond. In order to calculate the length and frequency of ETEs, the boundaries of social groups needed to be determined. As GPS data became available, it became clear that an earlier bait-marking study of the area did not accurately correspond to territory boundaries in this population (Fig. S2). Badgers reach maximum ranging in summer (Kowalczyk *et al*. 2000, 2006; Revilla and Palomares 2002; Woodroffe *et al*. 2016). Therefore, in areas where social groups are contiguous, as they are in Ireland, the shape of their summer ranges should reflect the physical borders between social groups. For each social group in each year, 95% MCPs were calculated using the summer GPS locations, *i.e.* June, July and August, of all the collared members of the social group (summer MCPs). Where we did not have data for these months, we included all available GPS data for social group members in that calendar year.

In order to map more realistic territory boundaries than MCPs provide, summer MCPs were plotted in ArcMap along with all of the GPS locations. A ‘geographical territory boundary’ was digitised based on the location of the summer MCP polygon, the real linear landscape features in the study area (roads, hedgerows, rivers and observed badger-paths) and the GPS locations of the relevant social group (Text S1). The boundaries for each year were compared and only changed if it was very clear that members of the social group were ranging differently from the years before/after. Most social groups were relatively stable. However, there were some examples of fission or fusion of social groups (Revilla and Palomares 2002; Macdonald *et al*. 2004), and the construction of the M11 altered the position of some territory boundaries adjacent to the road (Fig. S3).

To determine ETE distances, the ‘Generate Near Table’ tool in ArcMap was used to calculate the distance (m) between each of a badger’s GPS points and the nearest edge of the polygon representing the boundary of their social group in that year. All points that fell outside the polygon could be considered ETEs. However, as the Followit GPS collar locations can be inaccurate by up to 15m (Mullen *et al*. 2015), only locations that were >15m from the polygon where identified as ETEs in this study. On a given night, a badger may have recorded more than one GPS location outside its territory boundary. In such cases, only the GPS location that was furthest from the badger’s territory boundary was retained in the dataset. The frequency of ETEs per month (fETE), described as a proportion between 0 and 1, was calculated by dividing the number of active nights in a month by the number of ETEs made in that month. Active nights were defined as nights in which the badger was above ground for long enough to record GPS locations and wore a working collar. The fETE dataset also included collared badgers that never went on ETEs.

For each dataset, in addition to the response variables (ND, HR size, ETE distance and fETE), the following data were included: badger ID, social group ID, sex, age cohort, month, year, roadworks phase and roadworks adjacency. The data management process resulted in an ND dataset of 18,954 records, a HR dataset of 890 records, an ETE distance dataset of 3,726 records and a fETE dataset of 892 records.

### Statistical analyses

Generalized linear mixed models (GLMMs) were used to analyse the effects of the road on ND, HR and ETE distance and frequency in R Version 3.4.0 (R Core Team 2016). To investigate the effect of the roadworks, roadwork phase (three levels: before; during; after) was included as a fixed factor. To investigate whether the roadworks impacted the behaviour of badgers living in social groups adjacent to the roadworks to a greater extent than those who were living further away during different construction phases, an interaction between roadworks phase and a second fixed factor, adjacency (two levels: yes; no), was also included in the model. Sex (two factor levels: male; female), age cohort (four factor levels: cub; young adult; older adult; aged adult) and calendar month were included as fixed factors to control for the effect these variables had on ranging behaviour. Badger ID, nested within social group ID, and year (un-nested) were specified as random factors. Where response variables were nonnormal, BoxCox transformations (Box and Cox 1964) were used to normalise the data and the Gaussian family was specified in the GLMM. Where transformation failed to normalise the data, or where it was inappropriate to normalise the data, alternative appropriate distribution families were specified in the GLMMs (Table S2).

GLMMs were fitted using the package *lme4* (Bates *et al*. 2015) and *glmmTMB* (Brooks *et al*. 2017). Model selection was conducted using the package *MuMIn* (Barton 2017). Model selection was conducted using an information theoretical approach (Burnham and Anderson 2003; Bolker *et al*. 2009). A list of models using all possible combinations of predictor variables was ranked by Akaike Information Criteria for small sample sizes (AICc). Because we were interested in the relative importance of the predictor variables, the top model, rather than the average of a top model set was chosen (Cade 2015). The glht function in the package *multcomp* (Hothorn *et al*. 2008) was used to perform Tukey post hoc tests. Due to the skewed and variable nature of our data, results are presented in the text as medians and interquartile ranges (IQR) of individual medians, rather than overall means and standard deviations of the entire datasets, which can be found in Tables S3, S5, S7, and S9.

## RESULTS

The effects of the disturbance caused by road construction on the behaviour of the badgers was considered by looking at each parameter of ranging behaviour; nightly distance travelled, home range size, and the distance and frequency of extra-territorial ranging.

### Nightly Distance Travelled

On average, across the study period, badgers travelled a median distance of 657m a night (IQR: 458m-806). The majority of journeys, as determined using the 95^th^ percentile (Byrne *et al*. 2014), were below 2km (Table S3). The longest distance a badger travelled in a single night was 11.25km (Table S3). The model that best explained the variation in the nightly distance dataset retained the following explanatory variables; sex and age (in interaction with one another), roadworks, adjacency, and month (Table S4).

Before the roadworks commenced, badgers travelled a median distance of 651m (IQR: 382-777m, N = 7713) each night (Fig. 2a). During the road construction period, the median ND travelled by badgers increased slightly, but significantly, to 673m (IQR: 470-911m, N = 5428) (Table 1). Similarly, a further increase in median ND to 848m (IQR: 675-927, N = 5813) occurred after the new motorway opened (Table 1).

**Fig. 2.**
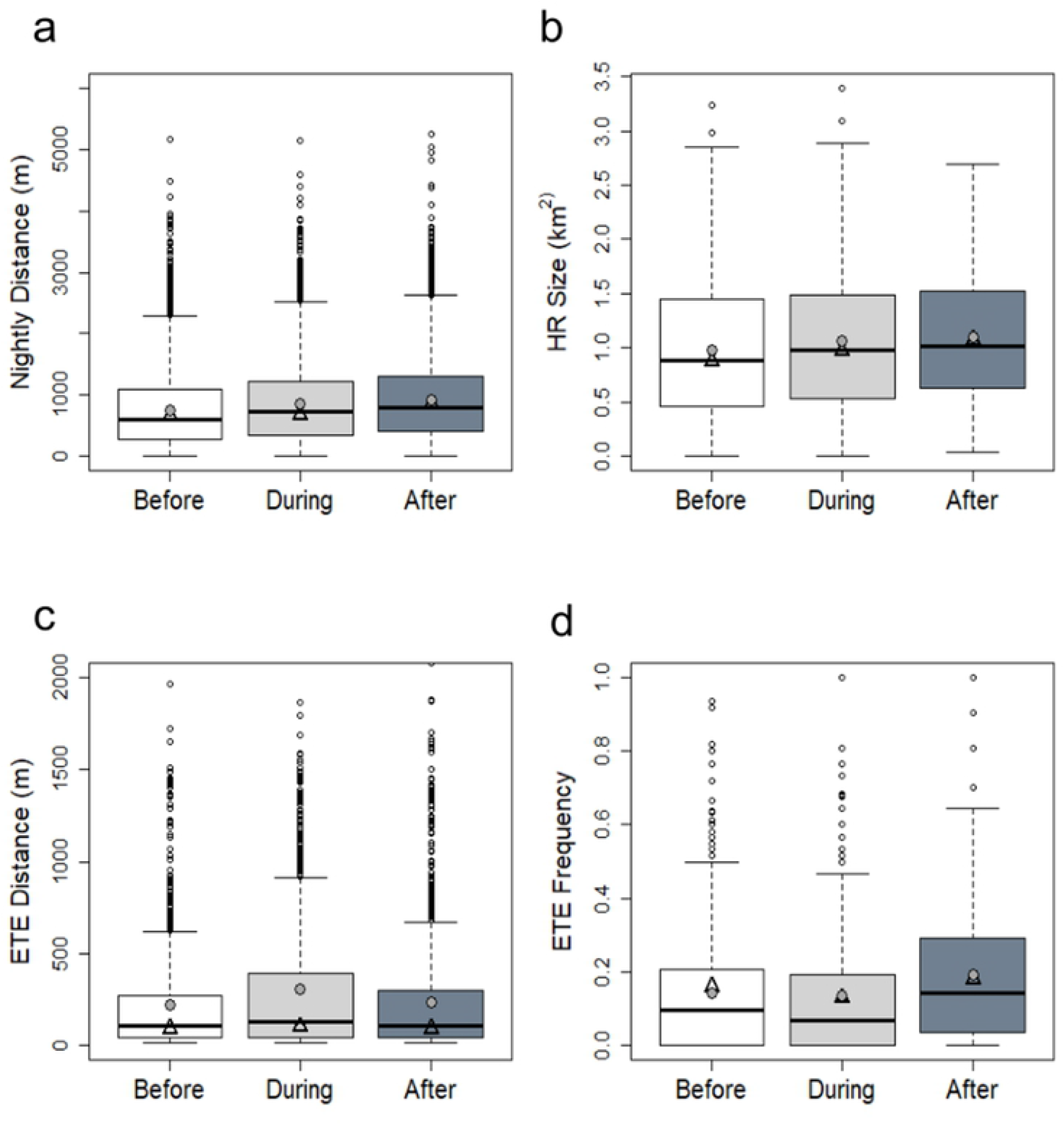
Boxplots for badger movement metrics in each phase of the roadworks. The before phase is represented in white, the during phase in light grey and the after phase in dark grey. Thick black bars represent cohort medians, filled grey circles represent cohort means and open black triangles represent the medians of individual medians. **a)** Nightly distance (m) by roadworks phase, y-axis limited to 6000m for clarity*; **b)** Home range size (km^2^) by roadworks phase; **c)** ETE distance (m) by roadworks phase, y-axis limited to 2000m for clarity* and **d)** ETE frequency (proportion between 0 and 1) by roadworks phase. * full graphs available in Fig. S4

**Table 1:** Results of Tukey *post hoc* tests for multiple comparisons of ND means for the different roadworks phases

| Comparison | Estimate | Std. Error | z value | Pr(>\|z\|) |  |
| --- | --- | --- | --- | --- | --- |
| Before-During | 0.054976 | 0.010482 | 5.245 | 3.13E-07 | *** |
| Before-After | 0.078302 | 0.012744 | 6.144 | 2.41E-09 | *** |
| During-After | 0.023326 | 0.008252 | 2.872 | 0.0047 | ** |

### Home range size

On average, across the study period, badgers had a monthly median HR size of 0.88km^2^ (IQR: 0.54-1.32km^2^). HR size ranged from 0.04km^2^ (40m^2^) to 3.4km^2^ (Table S5). We found no evidence for an effect of the roadworks on HR size, as neither roadworks phase nor adjacency to the road, either independently or in interaction with one another, were included in the model that best explained the data. The model that best explained the variation in home range size included sex and age (in interaction with one another) and month as explanatory variables (Table S6). Before the roadworks commenced, badgers had a median HR size of 0.87km^2^ (IQR: 0.49-1.38km^2^, N = 403) (Fig. 2b). During the roadworks, median HR size was 0.97km^2^ (IQR: 0.65-1.35km^2^, N = 260) while after the roadworks it was 1.07km^2^ (IQR: 0.68- 1.48km^2^, N = 227) (Fig. 2b).

### Distance of extra-territorial excursions

Of the 80 badgers in the study, 91% made ETEs (N = 73). At all times of year, over half of the collared badgers were making ETEs. Across the study period, the median straight-line distance that a badger travelled from its social group boundary was 99m (IQR: 61-167m). Most ETEs, as measured by the 95^th^ percentile, were less than 1km in length (Table S7). The maximum straight-line distance a badger travelled from its territory boundary was 4.2km (Table S7). We found no evidence that the roadworks influenced ETE distance, as neither roadworks phase nor adjacency to the road, either independently or in interaction with one another, were included in the best model. The model that best explained the variation in ETE distance included sex and age (in interaction with one another) and month as explanatory variables (Table S8).

Prior to the roadworks, the median distance of badger excursions was 90m (IQR: 65-127m, N = 1560) beyond their territory boundary (Fig. 2c). During the roadworks, the median distance of badger ETEs was 102m (IQR: 55-272m, N = 951) and after the new motorway was completed, the median ETE distance was 91m (IQR: 56-154, N = 1215) (Fig. 2c).

### Frequency of extra-territorial excursions

Of the badgers that were recorded leaving their territory, across the study period the median fETE was 0.14 (IQR: 0.10-0.23), which equates to 4.25 ETEs in a month. fETE ranged from 0 to 1, meaning that in a given month, some badgers went on ETEs every night, while others never left their territory (Table S9). Of these data, 83% (N = 742) were from badgers that were wearing functioning collars for the full month and 17% (N = 150) were from badgers whose collars functioned for only part of the month. We allowed for incomplete months by using the proportion of active nights. Only 6% (N = 54) of records were from partial months where the number of active nights was less than 2 weeks. We are therefore confident that our data are representative of typical ETE behaviour in this population.

The model that best explained the variation in fETE included sex, month and roadworks as explanatory variables (Table S10). Roadworks phase had a significant effect on the frequency with which badgers made ETEs (Table S10). Before the roadworks the median fETE was 0.16 (IQR: 0.10-0.23, N = 294). During the roadworks the median fETE was 0.13. The difference between the before and during phases was not significant (Table 2). However, badgers made ETEs significantly more frequently after the roadworks (median fETE 0.18, IQR: 0.10-0.25, N = 200) compared to the periods either before or during roadwork construction (Fig. 2d, Table 2). On average, in a given month, badgers made ETEs on 4.9 nights before the roadworks, on 3.9 nights during the roadworks, and on 5.5 nights after the roadworks.

**Table 2.** Results of Tukey post hoc tests for multiple comparisons of fETE means for the different roadworks phases

| Comparison | Estimate | Std. Error | z value | Pr(>\|z\|) |  |
| --- | --- | --- | --- | --- | --- |
| During – Before | 0.01907 | 0.02189 | 0.871 | 0.3837 |  |
| After – Before | 0.07418 | 0.0277 | 2.678 | 0.0198 | * |
| After – During | 0.0551 | 0.02028 | 2.717 | 0.0198 | * |

The period before the roadworks was much longer (41 months) than either the ‘during’ or ‘after’ phases (22 and 14 months respectively). In order to assess the biological significance of any changes in ranging behaviour caused by the roadworks, it is useful to consider how variable the badgers’ ranging was between years before any disturbance. We therefore considered the yearly medians, *i.e.* the median value for each year, for each of the movement metrics in the period before the roadworks commenced (Table 3). This helps to place any changes during and after the roadworks into the context of natural variation.

**Table 3.** Summary of median nightly distance (ND), home range (HR) size, extra-territorial excursion (ETE) distance and frequency (fETE) during each phase of the roadworks. The variation in annual median in the ‘before’ phase is included in the second row for each movement metric. The statistical significance of differences between the ‘during’ and ‘after’ phases when compared with the ‘before’ phase are marked with asterisks (*** < 0.001, * < 0.05), while (ns) signifies no significant difference.

|  | Median ND<br>(m) | Median HR size<br>(km<sup>2</sup>) | Median ETE<br>(m) | Median fETE |
| --- | --- | --- | --- | --- |
| Before roadworks | 651 | 0.87 | 90 | 0.16 |
| Variation in annual median before roadworks | 551-640 | 0.72-1.13 | 90-142 | 0.13-0.16 |
| During roadworks | 673 *** | 0.97 (ns) | 102 (ns) | 0.13 (ns) |
| After roadworks | 848 *** | 1.07 (ns) | 91 (ns) | 0.18 * |

### How often did badgers cross the N11/M11 before, during and after the roadworks?

In addition to the above movement metrics, we also calculated the number of times that badgers were recorded crossing the N11 road before and during the roadworks, and the M11 after the roadworks (Table S11). Badgers crossed the N11 road 75 times before the roadworks (1.8 crossing/month) and 22 times during the roadworks (1.7 crossings/month). After the roadworks badgers crossed the M11 motorway, using underpasses, 140 times (10 crossings/month). The badgers that had high levels of road crossing activity in the ‘after’ phase *i.e.*, >3 crossings/month came from just three social groups (outlined in Fig. 3 a-d). The majority were badgers from two of these social groups that were using underpasses to maintain access to parts of their territories (The Driving Range and The Briars) that would otherwise have been cut off by the new motorway. Badgers from a third social group (Hawthorn) were using an underpass to make ETEs to a neighbouring territory.

**Fig. 3.**
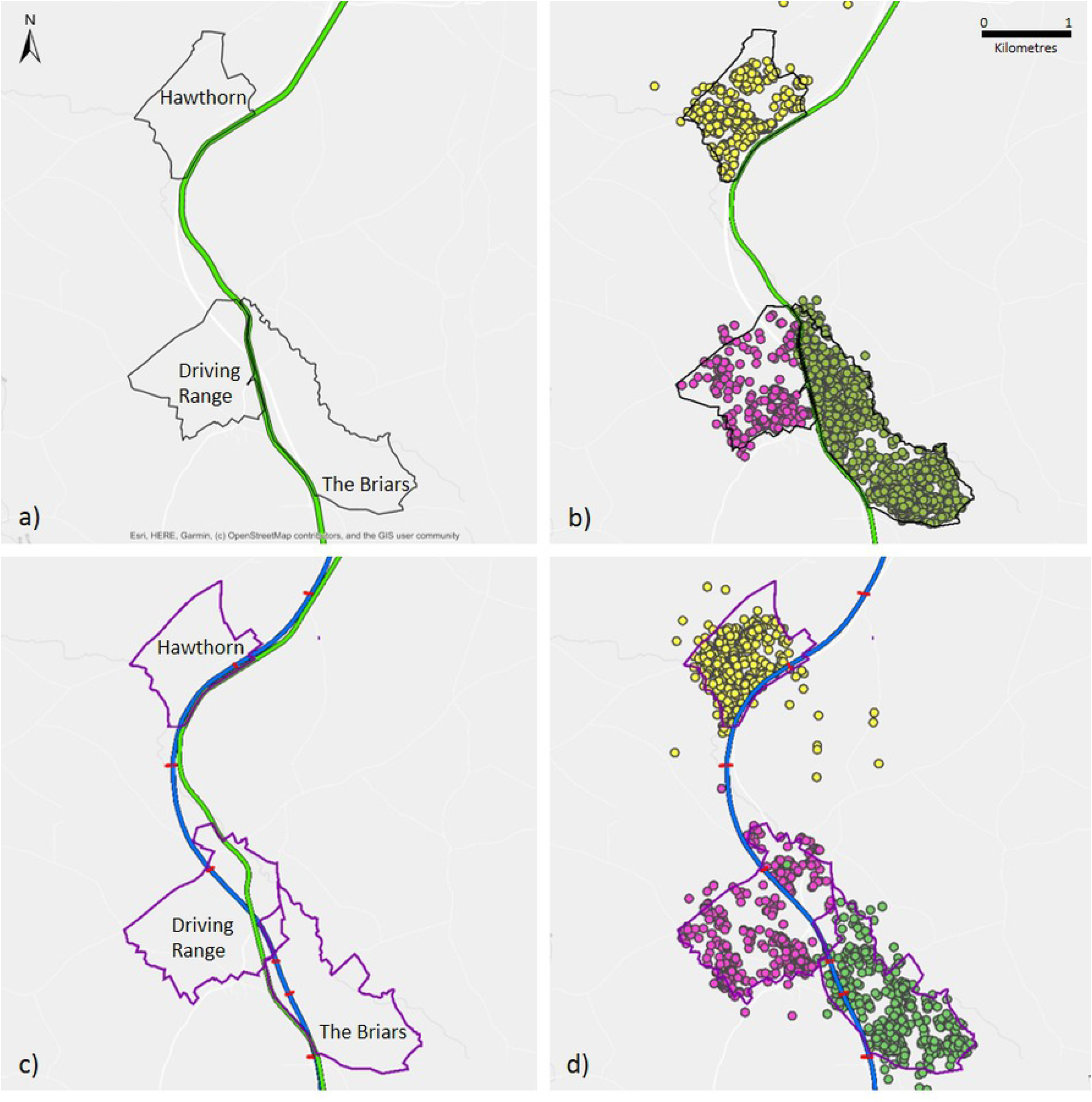
Territory Boundaries and Road Crossings. **a)** Territory boundaries of three of the social groups before the roadworks outlined in black; **b)** GPS locations for badgers from each of these social groups before the roadworks: Hawthorn in yellow, Driving Range in pink and The Briars in green; **c)** territory boundaries of the same three social groups after the roadworks outlined in purple and **d)** GPS locations for badgers from of these each social groups after the roadworks. The location of the N11 road is represented by a green line. The location of the M11 motorway is represented by a blue line. The locations of underpasses are represented by red dashes intersecting the M11. The N11 has been omitted from map d) for clarity.

## DISCUSSION

The disturbance caused by the process of road construction has the potential to impact the ranging behaviour of animals (Kohn *et al*. 1999; Klar *et al*. 2009; Lesmerises *et al*. 2013). Increased movement of badgers between social groups has been associated with an increase in the prevalence of TB in those social groups (Rogers *et al*. 1998; Riordan *et al*. 2011). However, the road upgrade considered here had very little effect on the ranging behaviour of badgers in the study area. The period before the roadworks spanned nearly 3.5 years and therefore acted as a record of normal behaviour. In considering any changes in the four movement metrics, it is helpful to bear in mind the natural variation between years as this gives a context for the values obtained during and after the roadworks (Table 3).

### Nightly Distance Travelled

The median distance moved by badgers in a night increased by 197m in total over the study period, by 22m a night during the construction phase, and by another 175m a night after the construction phase. This increase in ND suggests that the badgers were disturbed to some extent by the construction process and changes to the surrounding landscape. The fact that ND did not return to pre-roadworks distances once road construction was completed could indicate that the disturbance caused a permanent change in ranging behaviour, or that a monitoring period of greater than 12 months is required to capture a return to normal behaviour. While the median ND in the ‘before’ phase was 651m, there was a lot of variation in NDs across these years (Table 3). The values obtained during and after the roadworks fall outside this normal variation, suggesting a biologically significant effect on the badgers’ behaviour. However, it should be noted that the maximum recorded ND was 11.25km (Table S3), so an extra 1.75% (197m) in a night would represent a small proportion of the total distance the badgers were able to travel (11.25km).

It is also likely that the mitigation measures on the motorway (continuous badger-proof fencing and underpasses) had an effect on ND in the ‘after’ phase. Although an interaction between roadworks and adjacency was not included in the top model, badgers living in social groups located next to the N11/M11 may have had to travel further within their own territories once road construction was completed to maintain access to all areas of their territory (Fig. 3d), since a further increase in ND was seen after road construction was completed. Indeed, ND of badgers living in social groups adjacent to the N11/M11 increased by on average 168m during road construction, and by a further 68m after the roadworks, a total of 237m. While ND also increased for badgers living in territories that were not adjacent to the N11/M11, the total increase for these badgers was only 78m on average (2m during and 78m after the roadworks).

### Home Range Size

The small increase in ND did not manifest in a corresponding increase in HR size, suggesting that any extra ranging was confined primarily to within territory boundaries, indicating that territoriality in the area was not disrupted. This interpretation was strengthened by the finding that the median HR size both during and after the roadworks fell within the yearly range of median HR sizes recorded in the ‘before’ phase (Table 3). While there was no significant change in mean HR size of badgers, there was an inevitable alteration to the boundaries of the social groups immediately adjacent to the roadworks. Some social groups lost access and others gained access to land previously belonging to a different social group, however these changes were relatively small (Fig. S3). Potential loss of territory was mitigated for some social groups by the location of underpasses (Figs 3c and 3d) that facilitated normal ranging behaviour.

### Extra-Territorial Excursions

A possible consequence of an increase in ND might be an increase in the distance of ETEs. However, there was no change in ETE distance across the study period (Table 3). In a situation where disturbance leads to a breakdown in territoriality, ETE distance might be expected to increase. However, even though badgers were moving around more within their own territories during and after road construction, they remained as cautious about how far they trespassed into other territories as they had been before the roadworks commenced. This suggests that territory boundaries continued to be marked and defended as effectively as before the disturbance.

Similarly, any reduction in territoriality might be expected to lead to an increase in the frequency with which badgers undertook ETEs. Before the roadworks, badgers made, on average, 4.9 ETEs in a given month. However, there was no significant change in the frequency of ETEs during the roadworks. Once construction had finished and the new motorway was opened, the frequency of ETEs did increase, albeit by less than one additional ETE per month, to on average 5.5 ETEs a month. On average, ETEs were approximately 100m in length, and the majority were less than 1km. The average distance between main setts in the study area was 1.3km. The majority of ETEs were therefore likely to be into adjacent social groups. Our findings suggest that although the frequency of ETEs increased, the potential for badgers to interact with animals with which they had not interacted prior to the roadworks did not increase, since there was no increase in ETE distance. The availability of underpasses may have facilitated safe and successful ETEs (FIG. 3d) across a major road, a journey that was previously very risky.

### Badger Mortality on the New Motorway

A reduction in population density due to culling or persecution has been found to disrupt ranging behaviour in badgers (Sadlier and Montgomery 2004; Sleeman and Mulcahy 2005; Carter *et al*. 2007). Roads are a major cause of mortality in badgers (Davies *et al*. 1987; Clarke *et al*. 1998; Dekker and Bekker 2010). The extensive mitigation measures used in this case have proved successful in preventing badger deaths on the new M11 motorway. There were only two badger deaths on the M11 in the 14-month post-construction study phase. These were due to problems with incomplete fencing. Between 2010 and 2016, *i.e.* during the course of the study, we know of 49 badgers that died in RTAs on the N11 (n = 28) and minor roads (n = 21) within our study area. This gives us an average mortality rate of 10-15% per year. This is an order of magnitude higher than has previously been reported for rural Ireland (Sleeman *et al*. 2012) and is similar to figures reported for Sweden (Seiler *et al*. 2004) but is half the mortality rate reported for Denmark (Dekker and Bekker 2010). We know that the death of a badger in our study area often prompted the movement of other badgers from different social groups into that group. If the new motorway had lacked effective fencing and underpasses, it is likely an increase in RTA-related mortality, along with associated movements between social groups would have occurred. Therefore, we recommend that the high level of mitigation measures used here is employed, and periodically checked, in all major road realignment and road building projects.

### Implications of these changes for TB transmission

Taken together, our results suggest that any effect of the roadworks on ranging behaviour in badgers was very small, and that territoriality was not disrupted in our population. While there was a small but significant increase in ND during road construction, this was not reflected in any corresponding increase of HR size or ETE distance. After the new motorway was completed a further increase in nightly distance, and an increase in the frequency of ETEs was seen. However, this was not accompanied by a corresponding increase in home range size or ETE distance, suggesting that additional interactions within the social network were not formed as a result. It is possible that during the course of the roadworks, badgers responded to the disturbance by consolidating their own territories by ranging more within them, possibly marking border latrines more frequently. Only after the roadworks had finished was there evidence that they then began to investigate potential impacts upon adjacent territories through more frequent ETEs, although they did not venture further afield.

From commencement of the study we vaccinated all captured badgers against TB, using Bacille Calmette-Guérin (BCG) vaccine (MacWhite *et al*. 2013). At the start of the study, based on blood testing and bacteriology, TB prevalence in badgers in the study area was 19%. All captured study area badgers have been vaccinated with BCG vaccine since April 2010 and *M. bovis* has not been isolated from any badgers since December 2013. Since then there have been 7 more capture events, and 270 samples submitted for culture. Inside the study area there have only been two herd breakdowns since April 2010 - one high risk breakdown (c.15 reactor animals) on the edge of the study area in 2012, and one low risk breakdown (1 reactor animal) in the middle of the study area in March 2014. Both breakdowns were attributed to badgers. Since 2014, there have been no further breakdowns within the study area. In 2015, there was a complete herd depopulation in a farm directly adjacent to the study area. However, this was conclusively demonstrated to be due to an infected cow being brought into the herd. Vaccination of badgers mitigated against the possible transmission of TB between badgers both through bite wounding, and through activation of latent infections due to stress associated with the roadworks (Gallagher and Clifton-Hadley, 2000; Chambers et al. 2010; George *et al*. 2014).

Due to the fact that we vaccinated the badgers in our study area, we could not investigate whether there were any changes in TB infections of local cattle herds directly attributable to a change in badger ranging behaviour, that is, a potential ‘perturbation’ effect (Cheeseman *et al*. 1993; Tuyttens *et al*. 2000a, b; Donnelly *et al*. 2006, 2007; Carter *et al*. 2007; Jenkins *et al*. 2008). Badger movements into and out of neighbouring social groups are associated with increased prevalence of TB in those groups (Rogers *et al*. 1998; Riordan *et al*. 2011). Given the small degree of disruption to ranging behaviour evidenced, we believe it is unlikely that any increase in TB breakdowns in cattle would have occurred as a result.

It must be noted that the findings herein can only be extrapolated to other projects involving an upgrade of an existing road that is likely to act as a territory boundary (Clarke *et al*. 1998; Frantz *et al*. 2010). In greenfield sites, *i.e.* the construction of a new road over agricultural land, and potentially through the middle of existing, unvaccinated social groups, we cannot be certain whether ranging behaviour would be disturbed, nor whether there would be an associated perturbation effect on bTB in the area. However, the lack of disturbance seen, particularly in those social groups which lost territory sections in this road realignment, lead us to suggest that, where effective mitigation is provided, there is no expectation of a perturbation effect arising from road construction. We recommend that mitigation in the form of continuous badger-proof fencing be placed along the entire length of all major new road-builds and upgrades through areas occupied by badgers, and that underpasses should be located in appropriate places. This should prevent RTAs and minimise disturbance to the ranging behaviour of the badgers. We also recommend BCG vaccination of badgers in advance of major earthworks.

### Conclusion

The objective of this project was to ascertain the effects of a major road upgrade and realignment on the ranging behaviour of the badgers in the surrounding area. Road construction proved to be of minor consequence to the badgers’ movements or territoriality. Our results demonstrate that badgers can adapt to the considerable environmental disturbance resulting from major roadworks. Given the modest disruption in ranging behaviour seen in this badger population, both during and after road construction, we believe it is unlikely that a fully-mitigated major road upgrades would result in a perturbation effect sufficient to increase TB in the local cattle.

## ACKNOWLEDGEMENTS

We would like to thank Mark Foley and Denis Foley for their field work expertise, and all of the landowners for access to their lands.

